# Bridge-like lipid transfer protein family member 2 suppresses ciliogenesis

**DOI:** 10.1101/2023.12.07.570614

**Authors:** Jan Parolek, Christopher G. Burd

## Abstract

Bridge-like lipid transfer protein family member 2 (BLTP2) is an evolutionary conserved protein with unknown function(s). The absence of BLTP2 in *Drosophila melanogaster* results in impaired cellular secretion and larval death, while in mice (*Mus musculus*), it causes preweaning lethality. Structural predictions propose that BLTP2 belongs to the repeating β-groove domain-containing (also called the VPS13) protein family, forming a long tube with a hydrophobic core, suggesting that it operates as a lipid transfer protein (LTP). We establish *BLTP2* as a negative regulator of ciliogenesis in RPE-1 cells based on a strong genetic interaction with *WDR44*, a gene that also suppresses ciliogenesis. Like WDR44, BLTP2 localizes to membrane contact sites involving the endoplasmic reticulum and the tubular endosome network in HeLa cells and that BLTP2 depletion enhanced ciliogenesis by serum-fed RPE-1 cells, a condition where ciliogenesis is normally suppressed. This study establishes human BLTP2 as a putative lipid transfer protein acting between tubular endosomes and ER that regulates primary cilium biogenesis.

**Significance statement:** We show the involvement of an ER-localized bridge-like lipid transfer protein, BLTP2, in ciliogenesis and establish that BLTP2 is enriched at organelle-organelle membrane contact sites involving the endoplasmic reticulum (ER) and the tubular endosome network (TEN). These results implicate, for the first time, the involvement of bulk lipid transfer between the ER and TEN in regulating ciliogenesis.

## Introduction

Organelle-organelle membrane contact sites (MCS) are important hubs for the exchange of lipids and ions between organelles. To understand the physiological functions of MCSs, it is essential to identify the organelles involved, the protein networks that form the MCS between organelles and the physiological processes that require lipid transfer. The bridge-like lipid transport proteins (BLTP), also called VPS13-family proteins, have been proposed to facilitate bulk lipid transport between organelle membranes via a hydrophobic groove that run the length of the BLTP (Adlakha et al., 2022). The best characterized BLTP is ATG2/BLTP4 which transfers lipids from the endoplasmic reticulum (ER) to the autophagosome cup membrane which drives the growth of the nascent autophagosome (Osawa et al., 2019; Valverde et al., 2019). Many MCS proteins associate with the ER membrane by binding to the ER-localized integral membrane proteins VAPA and VAPB (VAPA/B), and to other organelles via organelle-specific proteins. The VAPA and VAPB proteins recognize FFAT sequence motifs located within many lipid transfer proteins, however BLTP1 and BLTP2 do not possess FFAT motifs (Hanna et al., 2023). Rather, they possess a *N*-terminal membrane spanning segment that localizes them to the ER membrane (Castro et al., 2022; Neuman et al., 2022). To date, human BLTP family proteins have been shown to be associated with mitochondria, peroxisomes, the Golgi apparatus and nascent autophagosomes. These observations suggest that BLTP family proteins transfer lipids between the ER and other membrane organelle. The focus of this study is human BLTP2.

Loss-of-function genetic studies of BLTP2 from diverse species indicate that it participates in diverse cellular and physiological processes (Neuman et al., 2022). In plants, BLTP2 is required for root tip (*Arabidopsis thaliana* and *Physcomitrella patens*) and pollen tube growth (*Zea mays*) (Benfey et al., 1993; Cheng & Bezanilla, 2021; Procissi et al., 2003; Xu & Dooner, 2006) and in *Drosophila melanogaster,* it is required for secretion and endosomal protein trafficking (Neuman & Bashirullah, 2018). Loss of BLTP2 in laboratory mice (*Mus musculus*) results in preweaning lethality in heterozygous animals (Groza et al., 2023). In humans, overexpression of BLTP2 has been associated with poor prognosis of cancer patients (Liu et al., 2014; Song et al., 2006; Zhong et al., 2018), yet there is no established disease related to mutations or absence of BLTP2.

In this study we investigated the localization and function of human BLTP2. Our results show that human BLTP2 localizes to the ER network and is concentrated in ER domains that are associated with components of the tubular endosomal network (TEN). We further show that a protein, WDR44, encoded by a gene that genetically interacts with *BLTP2*, is closely associated with BLTP2 at ER-TEN membrane contact sites. In agreement with a previously established role for WDR44 role in regulating ciliogenesis, we report that BLTP2 functions as an inhibitor of primary cilium biogenesis.

## Results and Discussion

### BLTP2 and WDR44 suppress ciliogenesis by RPE-1 cells

To begin to elucidate possible BLTP2 genetic and protein interaction networks, we consulted the Cancer Dependency Map Portal (version 23Q2+Score, Chronos) (Tsherniak et al., 2017) to identify genetic co-dependencies involving the *BLTP2* gene (**Fig. 1A**). BLTP2 and WDR44 have high mutual co-dependency (R = 0.61), suggesting a significant genetic interaction between these two genes, leading us to postulate that these proteins function in a common genetic pathway. WDR44 was previously established as a negative regulator of ciliogenesis in serum-fed RPE-1 cells (10% FCS), a condition where ciliogenesis is normally suppressed (Walia et al., 2019). In the RPE-1 cell line used in this study, we observed that 7.0±3.8% cells growing in serum-containing medium possessed a cilium and this proportion increased to 69.9±3.8% for cells switched to serum-free medium for 48 hours (**Fig. 1D, E**). Upon depletion of WDR44 using siRNA transfection (**Fig. 1C, D, E**), we observed that 36.7±5.9% cells produced a cilium when growing in serum-containing medium, thus confirming previous findings (Walia et al., 2019). To assess a potential role for BLTP2 in ciliogenesis, we depleted BLTP2 in RPE-1 cells (**Fig. 1B, D, E**) and determined the impact on ciliogenesis. Like WDR44-depleted cells growing in nutrient replete conditions, BLTP2 depletion also potentiated ciliogenesis in serum-containing medium with 27.6±7.9% cells having a cilium (**Fig. 1D**) while not affecting cilium biogenesis in serum starvation condition (**Fig. 1E**). Importantly, depletion of WDR44 or BLTP2 had no obvious impact on the appearance of the cilium, defined by immunostaining of two components of the primary cilium, Arl13b and acetylated tubulin (**Fig. 1E**) (Larkins et al., 2011). These results suggest that BLTP2 and WDR44 function in a regulatory pathway that suppresses ciliogenesis in RPE-1 cells.

**Figure 1:**
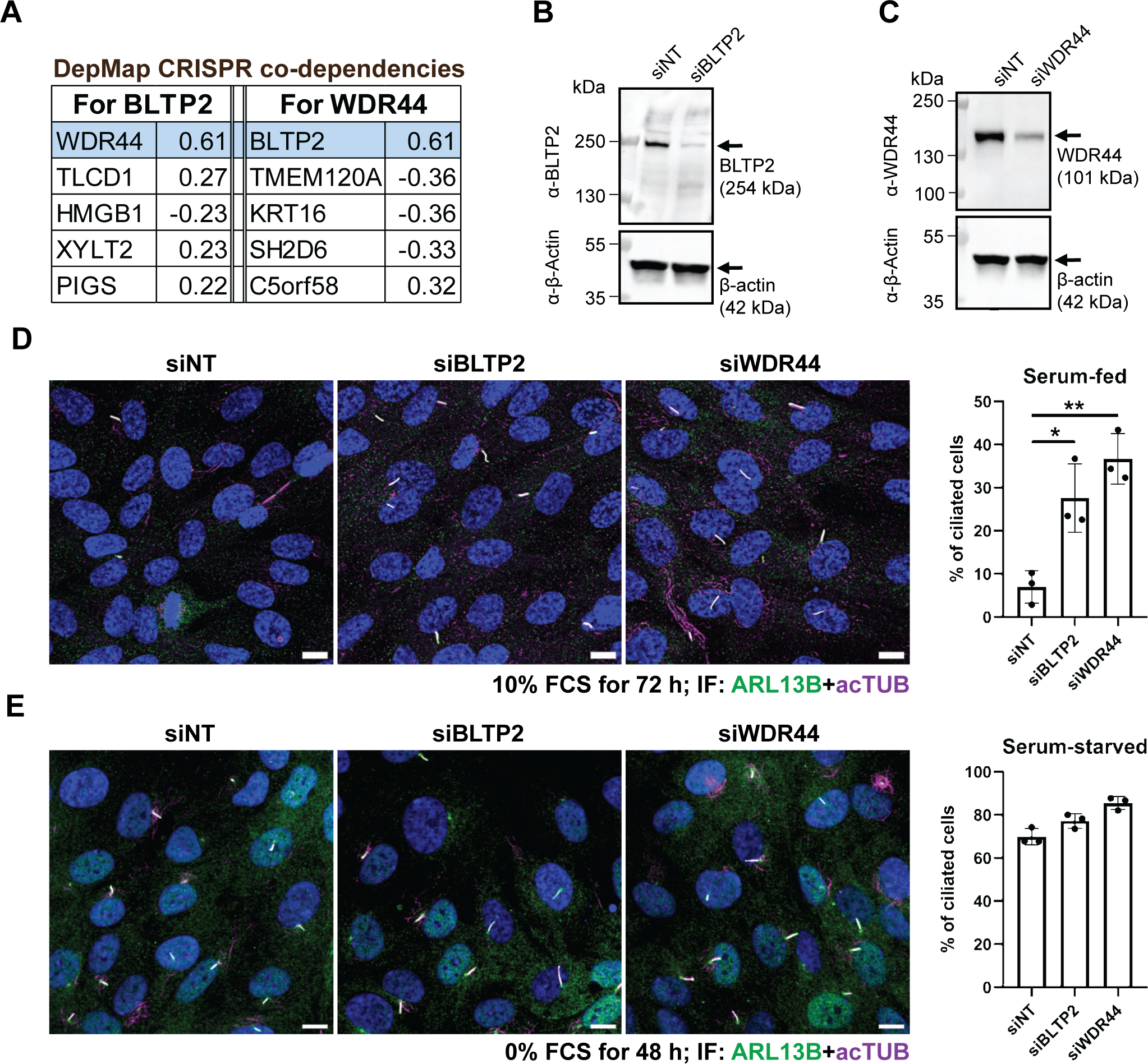
BLTP2, like WDR44, suppresses ciliogenesis in nutrient replete cells. **(A)** Top CRISPR co-dependencies show a significant genetic interaction between BLTP2 and WDR44. Source: DepMap Portal (version 23Q2+Score, Chronos). **(B)** WB showing BLTP2 depletion in siBLTP2-treated cells. **(C)** WB showing WDR44 depletion in siWDR44-treated cells. **(D)** RPE-1 cells were treated with siNT, siBLTP2 or siWDR44 and grown for 72 hours in the presence of 10% FCS. Cells were fixed and immunostained using α-Arl13b and α-acetyl-Tubulin antibodies (n>700 cells × 3). Scale bars: 10 µm. **(E)** RPE-1 cells were treated with siNT, siBLTP2 or siWDR44 and grown for 24 hours in the presence of 10% FCS. Then, cells were grown for 48 hours in the absence of serum to induce ciliogenesis. Cells were fixed and immunostained using α-Arl13b and α-acetyl-Tubulin antibodies (n>150 cells × 3). Scale bars: 10 µm.

### BLTP2 localizes to distinct domains of the ER

Published studies established that BLTP2 of *D. melanogaster*, *S. cerevisiae* and *P. patens* (called “hobbit”, Fmp27 and YPR117W, and “SABRE”, respectively) localize to the endoplasmic reticulum (ER) (Cheng & Bezanilla, 2021; Neuman & Bashirullah, 2018; Neuman et al., 2022; Toulmay et al., 2022). Since the *N*-terminus of human BLTP2 is predicted to serve as an ER membrane anchor, we first determined if human BLTP2 also localizes to the ER of RPE-1 and HeLa cell lines. We immunostained fixed HeLa cells using an antibody to BLTP2 and observed an enrichment of BLTP2 signal in linear structures (≥2 µm) that were not observed in two independent HeLa cell lines that do not express BLTP2 (BLTP2-KO#1, BLTP2-KO#2) (**Fig. 2A**). We verified the absence of BLTP2 in the HeLa cell knockout cells lines by immunoblotting using an α-BLTP2 antibody (**Fig. 2B**). Due to the prominent unspecific staining of the nucleus by this antibody (confirmed by immunostaining of BLTP2 knockout cell lines), we also determined the localization of transiently over-expressed human BLTP2 with a *C*-terminal HA-epitope tag, allowing us to probe BLTP2 localization more specifically in transfected cells. Using α-BLTP2 immunoblotting, we confirmed BLTP2-HA expression at the expected size (258 kDa) (**Fig. 2C**). In most of the transfected cells, BLTP2-HA localized throughout the ER network without any specific enrichment (**Fig. 2D**). However, BLTP2-HA also localized to tubular structures (≥2 µm) in 18±6.3% cells (n>150 cells in each of three independent experiments) (**Fig. 2E**). We noted that the enrichment of BLTP2 at ER sub-domains was most prevalent in cells with low relative expression of BLTP2-HA. We additionally performed live cell fluorescence microscopy of U2OS, and COS7 cells transfected with plasmids encoding BLTP2-SNAP, allowing visualization of BLTP2 in living cells upon addition of SNAP ligand, and GFP-Sec61B, an ER resident protein, and we observed an exclusive ER localization of BLTP2-SNAP (**Fig. S1A**). These results show that human BLTP2 protein localizes to the ER of multiple mammalian cell lines and that in HeLa cells it appears to be enriched in subdomains of the tubular ER network.

**Figure 2.**
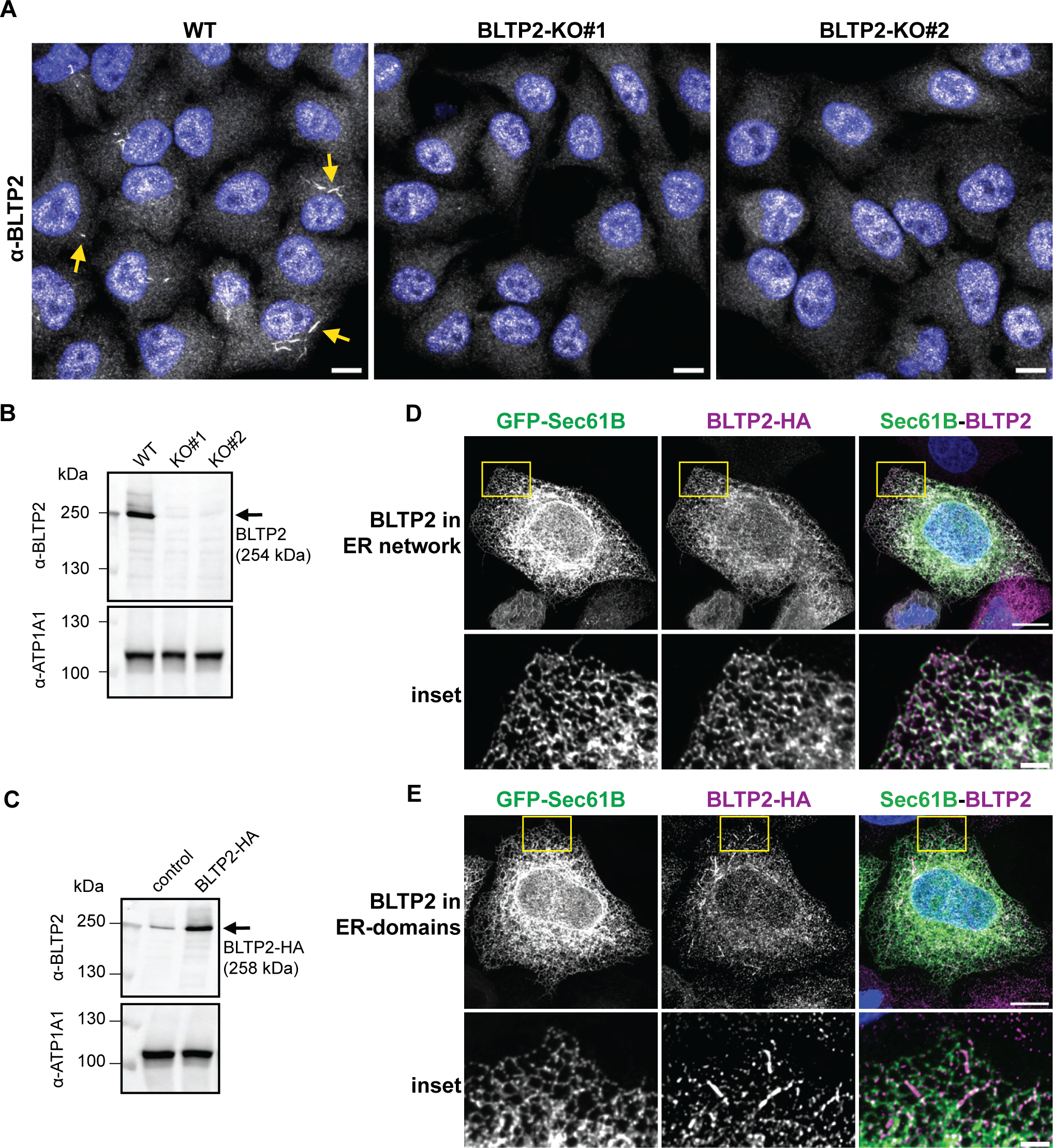
BLTP2 localizes to distinct domains of the ER network in HeLa cells. **(A)** BLTP2 is localized to pin structures that are present in WT and absent in KO cells. Example micrographs of HeLa WT and two BLTP2-KO cell lines that were fixed and immunostained using α-BLTP2 antibody. Scale bars: 10 µm. **(B)** A representative immunoblot shows BLTP2 absence in HeLa BLTP2-KO cell lines. Cell lysates were subjected to α-BLTP2 or α-ATP1A1 immunoblotting. **(C)** BLTP2-HA is expressed at expected size. A representative immunoblot shows the endogenous BLTP2 band (left) next to the BLTP2-HA band (right). The “control” lane contains lysate from non-transfected cells and the “BLTP2-HA” lane contains lysate from the cells transfected with BLTP2-HA. **(D, E)** BLTP2-HA localizes to pin structures in some cells (18±6.3%; n>150 cells × 3), while in most cells, it resides a general ER localization without any specific enrichments. Scale bars: 10 μm; scale bars of insets: 2 μm.

### BLTP2 and WDR44 are enriched at organelle membrane contact sites

In yeast cells, Fmp27 and YPR117W were observed to localize to ER-PM MCSs (Neuman et al., 2022; Toulmay et al., 2022). We observed that in HeLa cells, BLTP2 was not enriched at ER-PM MCSs, defined by the fluorescence pattern of GFP-MAPPER, a reporter of ER-PM MCSs (**Fig. S2**) (Chang et al., 2013). These data suggest that human BLTP2 does not localize to ER-PM contact sites in HeLa cells.

It was established previously that WDR44 possesses a FFAT (Phe-Phe-Ala-Thr) motif that binds to the VAPA/B ER integral membrane proteins which are common components of many MCSs involving the ER and other organelles (Baron et al., 2014) (**Fig. 3A**). In HeLa cells, WDR44 is enriched at a MCS involving the ER and the tubular endosomal network (TEN) (**Fig. 3A**) (Lucken-Ardjomande Hasler et al., 2020). The TEN of HeLa cells represents a subset of endosomes that are elongated (up to the length of the cell) tubules decorated with Rab8, Rab10 and other proteins (Etoh & Fukuda, 2019; Farmer et al., 2021; Giridharan et al., 2013; Lucken-Ardjomande Hasler et al., 2020). RPE-1 cells do not possess as elaborate TEN as that of HeLa cells, but other proteins involved in ciliogenesis (Rab8, EHD1, PACSIN2/Syndapin2, GRAF-1 and MICAL-L1) (Insinna et al., 2019; Vidal-Quadras et al., 2017; Xie et al., 2019) localize to the TEN of HeLa cells (Cai et al., 2014; Giridharan et al., 2013; Hattula et al., 2006; Sharma et al., 2009). To compare the localization of BLTP2 and WDR44 in RPE-1 cells, we transduced cells with a recombinant lentivirus that directs expression of BLTP2 fusion protein possessing a SNAP protein at its *C*-terminus, incubated with SNAP ligand, fixed and immunostained against WDR44 (**Fig. S3A, B**). In agreement with a published study, endogenous WDR44 localized to numerous puncta throughout the cytoplasm of RPE-1 cells, and we did not observe WDR44-coated tubules (**Fig. S3A**) (Lucken-Ardjomande Hasler et al., 2020). BLTP2 localized to the ER network (**Fig. S3A**). We next compared the localizations of BLTP2 and WDR44 in HeLa cells. Endogenous BLTP2 and transiently expressed HA-WDR44 co-localized on linear structures (>2 μm) that are similar in appearance to that of native BLTP2 (**Fig. 3B**). Importantly, BLTP2- and WDR44-structures are co-linear along segments of ER tubules, where BLTP2-SNAP is enriched (**Fig. 3C**).

**Figure 3:**
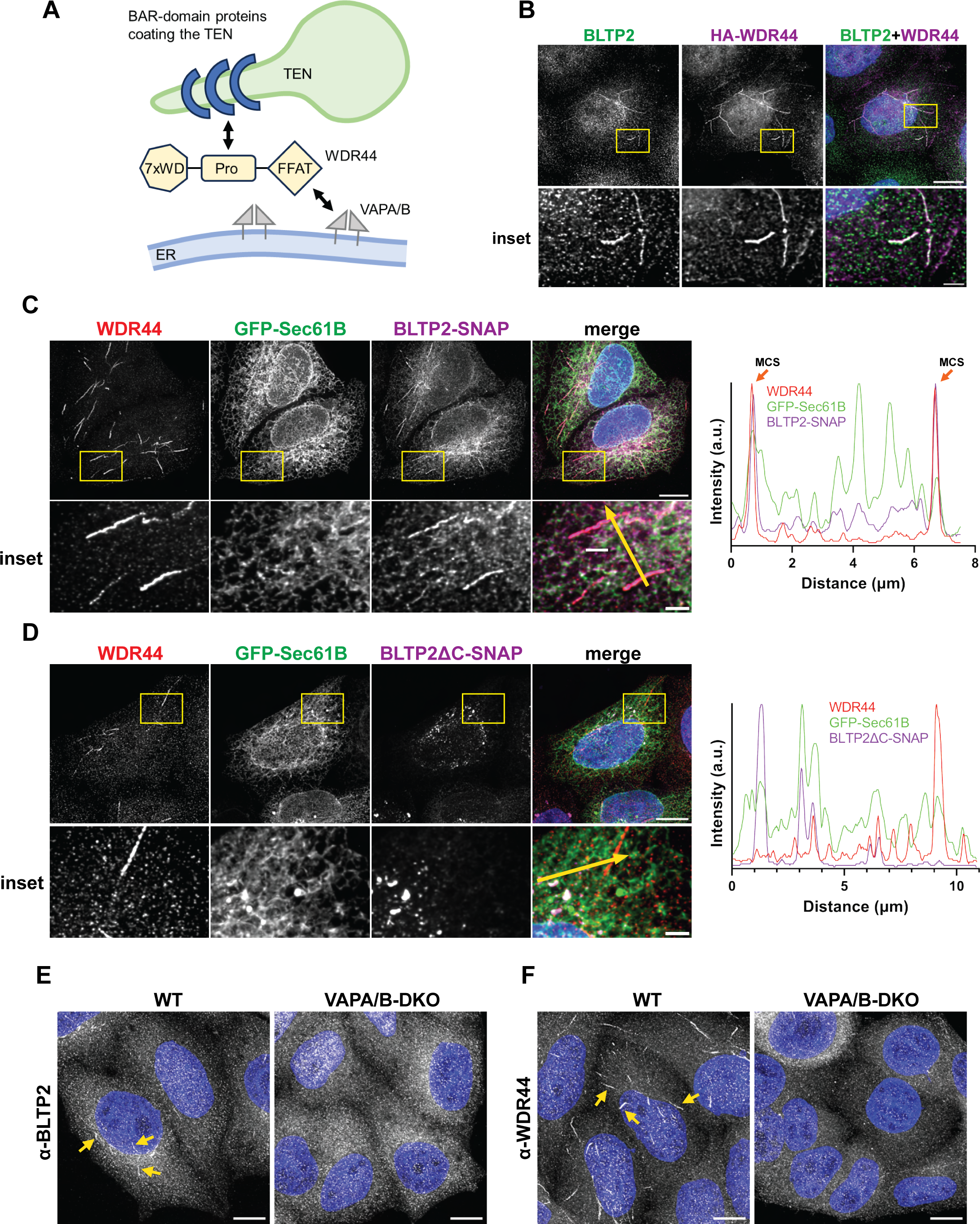
WDR44, a genetically interacting gene with BLTP2, co-localizes with BLTP2 to the tubular membrane network. **(A)** Schematic representation of WDR44, a protein connecting the ER and TEN. WDR44 interacts with SH3 motifs of BAR domain proteins on TEN by a polyproline tract (Pro) while the FFAT motif of WDR44 is recognized by VAPA/B in the ER. Adapted from Lucken-Ardjomande Häsler et al. (2020). **(B)** BLTP2 co-localizes with HA-WDR44 in HeLa cells. Example micrograph of a HeLa cell transfected with HA-WDR44 that was fixed and immunostained using α-BLTP2 and α-HA antibodies. Scale bars: 10 µm; scale bars of insets: 2 μm. **(C, D)** BLTP2 is enriched at the WDR44-positive tubules that are co-linear along the ER tubules, while the truncated mutant localizes to ER-puncta. Example micrographs of a HeLa cell transfected with GFP-Sec61B and BLTP2-SNAP or BLTP2ΔC-SNAP. Cells were first incubated with a SNAP-Cell ligand and then fixed and immunostained using α-WDR44 antibody. Line plot profiles are recorder from the path marked by the yellow arrows in the insets. Orange arrows mark the presence of membrane contact sites (MCS). Scale bars: 10 µm; scale bars of insets: 2 μm. **(E, F)** Tubule localization of BLTP2 and WDR44 is VAPA/B-dependent. HeLa WT and VAPA/B DKO cells were fixed and immunostained using α-BLTP2 or α-WDR44 antibody. Scale bars: 10 µm.

The *D. melanogaster* BLTP2 protein (2300 amino acids) was identified in a forward genetic screen for reduced body size and consequently called “*hobbit*” (Neuman & Bashirullah, 2018). The mutant *hobbit* phenotype is caused by nonsense mutations that delete *C*-terminal segments of the protein, resulting in deficiencies in secretion of glue granules and insulin-like polypeptides. One of the characterized *hobbit* alleles contained a deletion resulting in a *C*-terminal truncation (Q1882*) (Neuman & Bashirullah, 2018). To investigate the role of this *C*-terminal segment of human BLTP2 (2235 amino acids), we prepared a truncated version BLTP2(1-1814)-SNAP (BLTP2ΔC-SNAP) deletes the *C*-terminal 421 amino acids, corresponding to one of the original *D. melanogaster* loss-of-function alleles (**Fig. S1B**). We transfected HeLa cells with GFP-Sec61B to identify the ER and BLTP2(1-1814)-SNAP. The cells were incubated with SNAP ligand, fixed and immunostained against WDR44. The truncated version of BLTP2 (BLTP2(1-1814)-SNAP) localized to puncta within the ER network that were not associated with WDR44-positive TEN tubules (**Fig. 3D**). This result suggests that loss-of-function BLTP2 (i.e., *hobbit*) alleles that were obtained in *D. melanogaster* are deficient in association with the TEN, possibly because of accumulation in putative aggregates within the ER. Finally, in HeLa cells that do not express VAPA/B (Dong et al., 2016), WDR44 and BLTP2 were not enriched within ER domains (**Fig. 3E, F**). We conclude that association of ER-localized BLTP2 with the TEN is dependent upon the VAPA/B proteins and WDR44, which contains a FFAT motif (Baron et al., 2014).

### BLTP2 decreases proportion of cells with enlarged TEN

Next, we determined the impact of the loss of BLTP2 on WDR44 localization. In HeLa cells, WDR44 has been shown associated with sub-domains of the ER and TEN, that were similar in appearance of BLTP2 (**Fig. 2A**) (Lucken-Ardjomande Hasler et al., 2020). In BLTP2-KO cells, we observed WDR44 to extensively coat TEN membranes (**Fig. 4A**), likely by binding to the TEN coat protein, GRAF2 (Lucken-Ardjomande Hasler et al., 2020). Curiously, we observed that the proportion of cells possessing an elaborate TEN increased in absence of BLTP2 in the knockout HeLa cell lines (**Fig. 4B**). However, the area of WDR44 network (square pixels) per one cell with TEN remained unchanged (**Fig. 4C**). The phenotype of increased proportion of cells possessing WDR44 tubules was reverted by expression of BLTP2-HA (**Fig. 4D, E**). These results raise the possibility that BLTP2 is a negative regulator of the WDR44-associated TEN.

**Figure 4:**
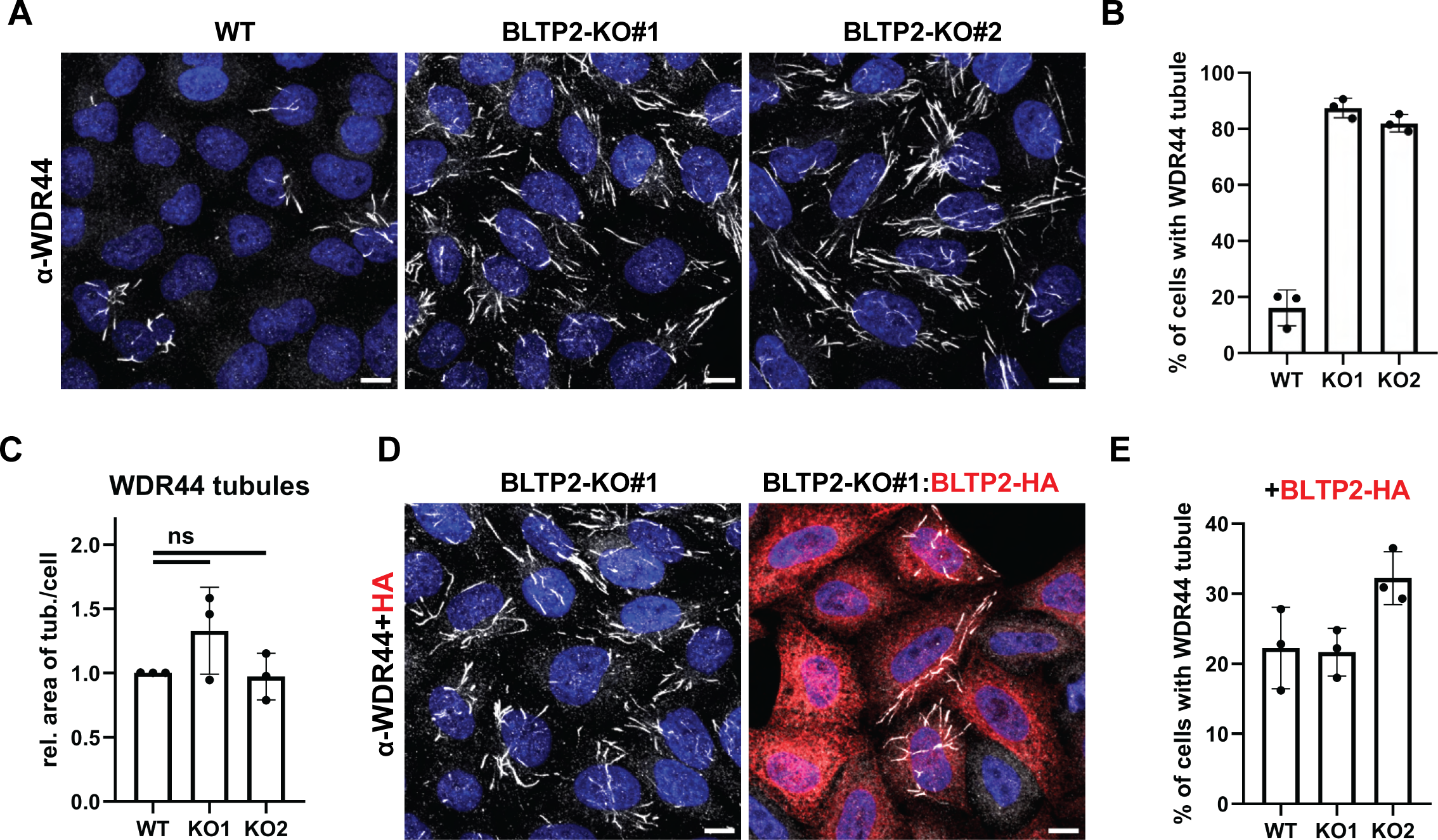
WDR44 network is present in higher proportion of cells lacking BLTP2. **(A)** HeLa WT and two BLTP2-KO cell lines were fixed and immunostained using α-WDR44 antibody. Scale bars: 10 µm. **(B)** Quantification of proportion of cells positive for enlarged WDR44 TEN for HeLa WT and two BLTP2-KO cell lines (n>150 cells × 3). **(C)** Quantification of relative area of WDR44 network per one cell positive for enlarged WDR44 TEN for HeLa WT and two BLTP2-KO cell lines (n>150 cells × 3). **(D)** Example micrograph of a HeLa BLTP2-KO#1 cells transfected with BLTP2-HA. Cells were fixed and immunostained using α-WDR44 and α-HA antibodies. Scale bars: 10 µm. **(E)** Quantification of proportion of cells positive for enlarged WDR44 TEN for HeLa WT and two BLTP2-KO cell lines transfected with BLTP2-HA (n>150 cells × 3).

### BLTP2 and WDR44 localize to tips of GFP-Rab8 and GFP-Rab10 coated tubules

In HeLa cells, the TEN can be decorated by GFP-Rab8 and GFP-Rab10 (Etoh & Fukuda, 2019; Lucken-Ardjomande Hasler et al., 2020). Rab8 was shown to be essential for endocytic recycling and ciliogenesis (Dhekne et al., 2018; Hattula et al., 2006; Nachury et al., 2007; Vidal-Quadras et al., 2017) and Rab10 has been implicated in diverse cellular processes, including basolateral protein sorting, macropinocytosis and, especially relevant for this study, ciliogenesis (Dhekne et al., 2018; Liu et al., 2020; Nakamura et al., 2020). To determine if BLTP2-associated tubules are part of the Rab8/Rab10 TEN, we transfected HeLa cells with plasmids encoding GFP-Rab8, GFP-Rab10 and GFP-Rab11, as WDR44 is an effector of Rab11, (Mammoto et al., 2000; Thibodeau et al., 2023; Zeng et al., 1999) and immunostained them to visualize BLTP2 or WDR44 (**Fig. 5A, B, D, E**). We did not observe GFP-Rab11 puncta to be associated with BLTP2- or WDR44 (**Fig. 5C, F**), as previously noted (Lucken-Ardjomande Hasler et al., 2020). We did, however, observe a striking enrichment of both BLTP2 and WDR44 at the ends of the GFP-Rab8- or GFP-Rab10-decorated tubules. With these results in mind, we suggest that the BLTP2-enriched domains of the ER correspond to ER-TEN MCSs and this enrichment at ER-TEN MCSs is required to suppress ciliogenesis in serum-fed RPE-1 cells.

**Figure 5.**
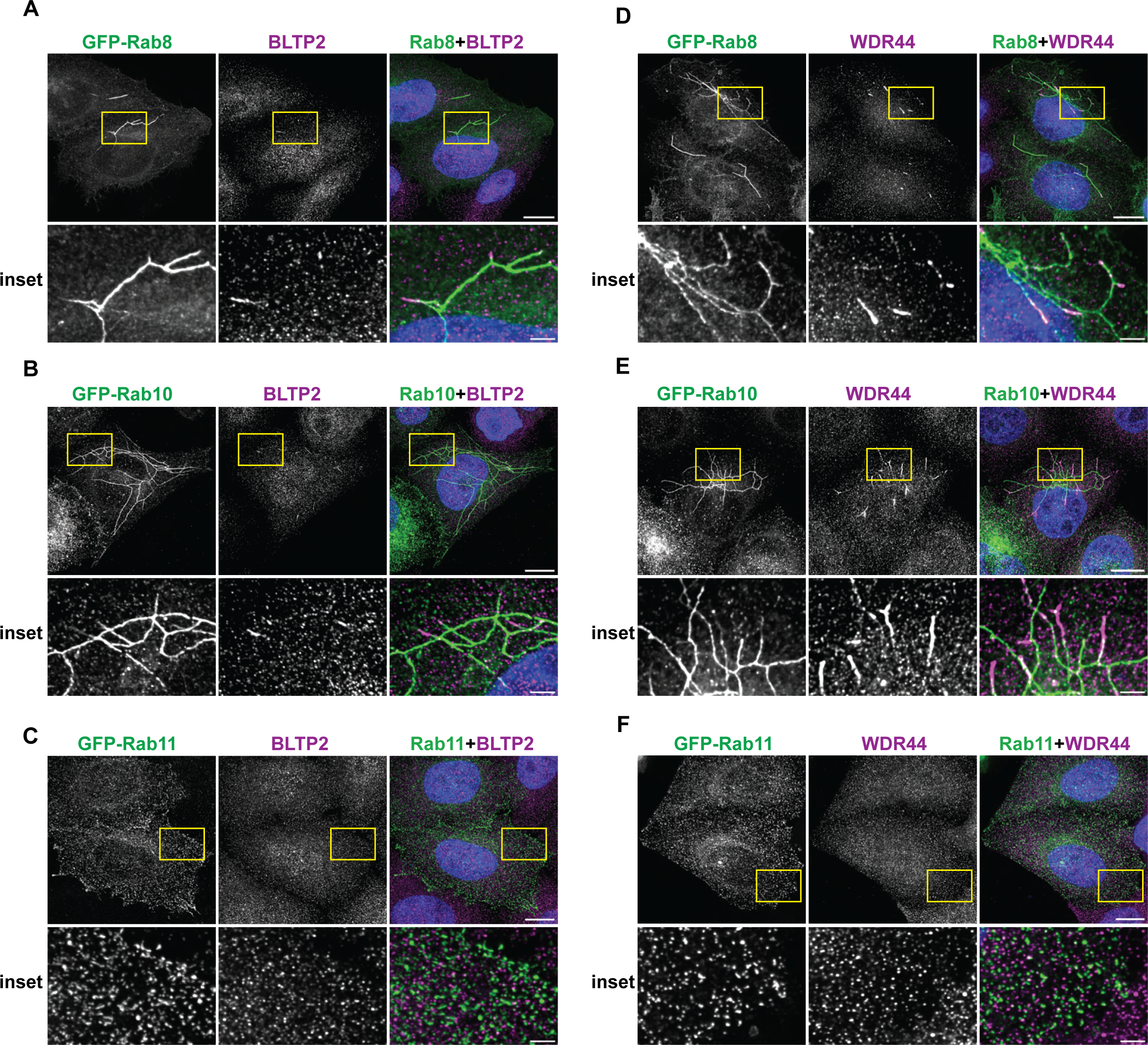
BLTP2 and WDR44 localize at tips of GFP-Rab8 and GFP-Rab10 tubular membrane network. **(A, B)** BLTP2 is enriched at TEN tips of GFP-Rab8 and GFP-Rab10. Transfected HeLa cells were fixed and immunostained using α-BLTP2 antibody. Scale bars: 10 µm; scale bars of insets: 2 μm. **(D, E)** WDR44 is enriched at TEN tips of GFP-Rab8 and GFP-Rab10. Transfected HeLa cells were fixed and immunostained using α-WDR44 antibody. Scale bars: 10 µm; scale bars of insets: 2 μm. **(C, F)** WDR44 and BLTP2 were not observed associated with GFP-Rab11. Transfected HeLa cells were fixed and immunostained using α-BLTP2 or α-WDR44 antibody. Scale bars: 10 µm; scale bars of insets: 2 μm.

The results of this study suggest that BLTP2 and WDR44 are common components of a pathway that suppresses ciliogenesis in serum-fed RPE-1 cells. In ciliated cells, a Rab11-to-Rab8 signaling cascade regulates vesicular trafficking from the endosome to the mother centriole, thereby contributing directly to the formation of the ciliary vesicle (Dhekne et al., 2018; Knodler et al., 2010; Lu et al., 2015; Nachury et al., 2007; Walia et al., 2019). WDR44 is an effector of Rab11^GTP^ in HeLa and RPE-1 cells and (Walia et al., 2019) proposed that the Rab11^GTP^-WDR44 complex suppresses ciliogenesis by sequestering Rab11^GTP^ from other Rab11^GTP^ effectors that promote ciliogenesis, including Rabin8 and FIP3 (Walia et al., 2019). Given the strong genetic interactions between the BLTP2 and WDR44 genes (Tsherniak et al., 2017) it is likely BLTP2 also regulates ciliogenesis by inhibiting the activities of pro-ciliogenesis factors indirectly (Walia et al., 2019). In HeLa cells, loss of BLTP2 increased the proportion of cells with an elaborate TEN, suggesting that it suppressed formation of the TEN. A relationship between regulation of ciliogenesis and formation of the TEN has not been reported previously. Although it has not yet been established if BLTP2 exhibits membrane lipid transfer activity, we note that a mutant form of human BLTP2, based upon a *D. melanogaster* loss-of-function *hobbit* allele, is not enriched at ER-TEN MCSs and therefore would be unable to mediate transfer lipid at these sites. Future studies will need to definitively determine if BLTP2 mediates lipid transfer between the ER and the endo-lysosomal network and, if so, how this activity influences signaling by Rab11, Rab8 and Rab10 signaling networks.

We consider it likely that BLTP2 and WDR44 are associated, directly or indirectly, at the ER-TEN MCS because BLTP2 and WDR44 co-localize at the tips of GFP-Rab8- and GFP-Rab10-decorated tubules in HeLa cells, and their mutual requirements to be enriched at these structures. Previous research using HeLa cells suggested a role for the TEN in plasma membrane recycling of CD59 and CD147 (Cai et al., 2011; Etoh & Fukuda, 2019) and anterograde trafficking of some neo-synthesized integral membrane proteins, including MMP14, E-cadherin and CFTRΔ508, from the endosome to the plasma membrane (Lucken-Ardjomande Hasler et al., 2020). We did not observe accumulation of the endocytosed CD59-antibody complex or neo-synthesized MMP14-GFP or E-cadherin-GFP in the TEN of BLTP2-depleted HeLa cells, so BLTP2 function is not required to support the export of these integral membrane proteins from the TEN. To the extent that these three proteins report all cargo export pathways from the TEN, the requirement for BLTP2 in regulating ciliogenesis does not, therefore, appear to be mediated by effects on vesicular trafficking to the plasma membrane via the TEN. Rather, BLTP2 may suppress ciliogenesis by altering the lipid dynamics and/or densities of unidentified integral membrane proteins that suppress ciliogenesis.

## Materials and Methods

### Cell culture and transfection

All HeLa cell-based assays utilized HeLa cells that were purchased from American Type Culture Collection and were a kind gift from Dr. Julia von Blume (Yale University). 293T cells originated from ATCC (CRL-3216) and were a kind gift from Dr. David Calderwood (Yale University). Cells were grown in DMEM (11965-092, Gibco) supplemented fetal bovine serum (FBS; 10% vol/vol; 16140-071, Gibco), unless otherwise noted. HeLa VAPA/B double knockout and companion parental cells were a kind gift from Dr. Pietro de Camilli (Yale University). hTERT RPE-1 cells originated from ATCC (CRL-4000) and were a kind gift from Dr. Derek Toomre (Yale University). RPE-1 cells were grown in DMEM/F12 (11330-032, Gibco) supplemented fetal bovine serum (FBS; 10% vol/vol; 16140-071, Gibco), unless otherwise noted. All cell lines were grown in humidified incubator at 37°C with 5% CO_2_ and were routinely screened for mycoplasma contamination using a mycoplasma detection kit (30-1012K, ATCC).

### Plasmids and siRNAs

Lipofectamine 2000 (11668-019, Invitrogen) was used for transient transfection of cells with plasmids in accordance with the manufacturer’s instructions (1 µg plasmid DNA with 2.5 µl of Lipofectamine 2000 in 1 ml). We noted that the formation of TEN could be perturbed by this transfection reagent, as transfected cells showed a decreased proportion of cells with WDR44 tubules. All analyses of transfected cells were done within 24 hours after transfection. Following plasmids were purchased from Addgene: GFP-MAPPER (#117721), GFP-Rab10 (#49472), pLenti-CMV-Neo-DEST(705-1) (#17392) and pSpCas9(BB)-2A-Puro (#62988). Gateway pDONR221 vector was purchased from Invitrogen (12536017). The pCMV5D_HA-WDR44 plasmid was purchased from MRC PPU Reagents and Services (DU26855) and a pFN21A-Halo-KIAA0100 plasmid (which served as a PCR template) was purchased from Promega (FHC00016). GFP-Rab8 and GFP-Rab11 were kind gifts from Dr. Shawn Ferguson (Yale University). The mEmerald-Sec61-C-18 and SNAP-Sec61B plasmids were kind gifts from Dr. Joerg Bewersdorf (Yale University) (Fuentes et al., 2023). pCMV-VSV-G envelope protein and pCMV-Δ8.91 viral packaging plasmids were kind gifts from Dr. David Calderwood (Yale University).

Oligonucleotides used for DNA sequencing and mutagenesis were synthesized by the Keck Biotechnology Resource Laboratory of Yale University. Cas9 plasmids targeting sequences *BLTP2* open reading frame were prepared by digesting pSpCas9(BB)-2A-Puro plasmid with BbsI (R3539S, NEB) and inserting duplexed primers caccGAGCCAAACACGGTTCCGGA and aaacTCCGGAACCGTGTTTGGCTC or caccGTGCAAGATCTGACGCCGAC and aaacGTCGGCGTCAGATCTTGCAC. To generate plasmids encoding epitope tagged BLTP2, human BLTP2/KIAA0100 ORF was amplified from pFN21A-Halo-KIAA0100 (FHC00016, Promega). An unstructured linker (sequence: GGSGGGGSGGGPGGGSGGSGGSGGG) was placed downstream BLTP2 ORF followed with 2xHA-tag (YPYDVPDYAGYPYDVPDYA) or SNAP-tag. BLTP2-SNAP(1-1814) was prepared from BLTP2-SNAP by PCR deletion of the relevant sequence. BLTP2-SNAP ORF was PCR-amplified using primers with attB1 and attB2 overhangs, GGGGACAAGTTTGTACAAAAAAGCAGGCTTCACCATGCCTCTGTTCTTCTCCGCGCTG and GGGGACCACTTTGTACAAGAAAGCTGGGTTTAACCCAGCCCAGGCTTGCCCAG, and using Gateway BP Clonase II Enzyme mix (11789020, Invitrogen) and pDONR221 vector, pDONR221_ BLTP2-SNAP was prepared. Then, using Gateway LR Clonase II Enzyme mix (11791020, Invitrogen) and pDONR221_BLTP2-SNAP, pLenti-CMV-Neo-DEST_BLTP2-SNAP was prepared. Similarly, SNAP-Sec61B ORF was PCR-amplified using primers with attB1 and attB2 overhangs, GGGGACAAGTTTGTACAAAAAAGCAGGCTATGGACAAAGACTGCGAAATGAAGC and GGGGACCACTTTGTACAAGAAAGCTGGGTCTACGAACGAGTGTACTTACCCCAAATGTG, and using Gateway BP Clonase II Enzyme mix (11789020, Invitrogen) and pDONR221 vector, pDONR221_SNAP-Sec61B was prepared. Then, using Gateway LR Clonase II Enzyme mix (11791020, Invitrogen) and pDONR221_SNAP-Sec61B, pLenti-CMV-Neo-DEST_SNAP-Sec61B was prepared. The sequences of all constructs were confirmed by DNA sequencing and are available upon request.

Lipofectamine RNAiMAX (13778-150, Invitrogen) was used for siRNA experiments according to the manufacturer’s instructions (30 pmol siRNA with 2.0 µl of Lipofectamine RNAiMAX in 1 ml). We noted that the formation of TEN could be perturbed by this transfection reagent, as transfected cells showed a decreased proportion of cells with WDR44 tubules. The effectiveness of siRNAs targeting BLTP2, and WDR44 was assayed by immunoblotting with an antibody raised against each protein. The following siRNA sequences were used: non-specific target (siNT), CGTTAATCGCGTATAATAC; BLTP2 (siBLTP2), GTCCTGATCAAGTGTGATAAATTTT, designed by IDTDNA (hs.Ri.KIAA0100.13.3); WDR44 (siWDR44), GTATAAGGGTTACGTCAAT (Walia et al., 2019). All siRNAs were purchased from Integrated DNA Technologies.

### Knockout cell line generation

The BLTP2-deleted cells were prepared by co-transfecting HeLa cells with two plasmids pSpCas9(BB)-2A-Puro targeting sequences GAGCCAAACACGGTTCCGGA and GTGCAAGATCTGACGCCGAC within the *BLTP2* gene. After transfection with Cas9 plasmids, HeLa cells underwent a 48-hour selection with 2 μg/ml puromycin (J67236.XF, Thermo Scientific Chemicals). The absence of BLTP2 in clonally selected cell lines was assayed by immunoblotting and immunofluorescence microscopy using an anti-BLTP2 antibody.

### Lentiviral production and transduction

HEK293T cells were co-transfected with pCMV-VSV-G, pCMV-Δ8.91 and pLenti-CMV-Neo-DEST_BLTP2-SNAP or pLenti-CMV-Neo-DEST_SNAP-Sec61B. The cell media was replaced 6 hours after transfection. Medium containing recombinant lentivirus was collected after 48 h later and passed through a low protein binding 0.45μm filter unit. hTERT-RPE-1 cells were infected with recombinant lentivirus-containing medium, supplemented with 10 µg/ml polybrene (TR-1003-G, Sigma). The next day, the medium was replaced, and cells were further grown in medium supplemented with 1000 µg/ml G418 (11811-031, Gibco).

### Ciliogenesis assay

For ciliogenesis assays, hTERT-RPE-1 cells were plated in 6-well plates with 12mm coverslips and incubated overnight. For serum starvation assays 400,000 cells were seeded on a 3.5cm dish and for the nutrient replete condition, 120 000 cells were seeded. The next day the cells were transfected with siRNA and incubated for 16 hours. Afterwards, cells were either incubated with DMEM/F12 containing 10% FBS (normal medium) or 0% FBS (starvation media) and incubated for additional 48 hours in media that was replaced daily. After 72 hours from siRNA treatment, cells were either lysed and used for immunoblotting or fixed and used for immunostaining. For ciliogenesis quantification using fluorescence microscopy, only Arl13B+acetylated Tubulin double-positive cilia were included in the analysis.

### Fluorescence microscopy and live cell imaging

The fluorescence micrographs shown in all figures are representative of at least three independent experiments. The micrographs were captured using a SoRa CSU-W1 (Yokogama) spinning disk confocal workstation (Nikon) based on an inverted microscope (Nikon Ti2-E) and ORCA-FusionBT back-thinned camera (Hamamatsu). Images were captured using 60× oil lens (N.A. 1.4), 30CC immersion oil F and NIS-Elements Advanced Research Package (Nikon). All presented micrographs are maximum intensity projections of 0.2µm confocal slices covering the whole volume of cells. Images were denoised and deconvoluted using NIS-Elements Advanced Research Package (Nikon). The images displayed in the figures were adjusted in brightness/contrast using Fiji (Schindelin et al., 2012).

For immunostaining, cells were cultured on glass circular coverslips (72230-01, EMS), fixed for 10 minutes with 4% PFA and permeabilized and blocked in 0.2% saponin and 1% BSA in PBS (SBPBS). Samples were incubated with primary antibodies diluted in SBPBS for 60 minutes and then with secondary antibodies in SBPBS (with 0.5 µg/ml Hoechst 33258) for 45 minutes. Primary antibodies used in this study include anti-acetyl-α-tubulin (IF 1:500; 66200-1-Ig, Proteintech), anti-ARL13B (1:500; 17711-1-AP; Proteintech), anti-BLTP2/KIAA0100 (1:250; HPA042781; Sigma), anti-HA (1:500; G036; abm), and anti-WDR44 (1:500; A301-440A, Bethyl). Fluorescent secondary antibodies were purchased from Invitrogen. Coverslips with immunostained cells were attached to glass slides using ProLong Glass Antifade Mountant (P36980, Invitrogen). Cells were imaged using the SoRa spinning disk confocal workstation described above.

For live cell imaging, cells cultured in glass bottom dishes (P35G-1.5-14-C, Mattek) were transfected and incubated with 0.5 µM 647-SiR SNAP ligand (S9102S, NEB) and 0.5 µg/ml Hoechst 34580 in DMEM (11965-092, Gibco) containing FBS (10% vol/vol; 16140-071, Gibco) for 30 minutes. Cells were imaged in 0.5 µg/ml Hoechst 34580 in DMEM containing 10% FBS at 37°C with 5% CO_2_.

### Immunoblotting

Post-nuclear supernatants (PNS) of cells were made as follows. Cells were washed in PBS, resuspended in lysis buffer containing 20 mM Tris-HCl pH 7.5, 1 mM EDTA, 150 mM NaCl, 1% Triton X-100 and protease inhibitor cocktails (04693116001, Sigma), briefly sonicated and then centrifuged for 1 min at 16,000 × g. The protein content of PNS samples was measured by Coomassie-protein quantification solution (23200, Pierce) and normalized by dilution in lysis buffer. PNS samples were mixed with 5× SDS-PAGE sample buffer containing 250 mM Tris-HCl pH 6.8, 50% glycerol, 10% sodium dodecyl sulfate, 10% β-mercaptoethanol, 0.025% bromophenol blue and incubated in room temperature for 10 minutes. Proteins were resolved by SDS-PAGE (4568026, BioRad) and then transferred to nitrocellulose membranes (1620115, BioRad). Bulk proteins present on the membrane were visualized by Ponceau S staining (0.1% Ponceau S in 5% acetic acid) and then the membranes were blocked for 60 minutes by incubation in 5% non-fat milk (AB10109-01000, AmericanBio) in 0.2% Tween 20 in PBS (PBST). Membranes were incubated in primary antibodies at the indicated dilutions in PBST at 4 °C for 16 hours. The antibodies used in this study include anti-β-actin (WB 1:1000; 3700S, CST), anti-ATP1A1 (WB 1:5000; ab76020, abcam), anti-BLTP2/KIAA0100 (WB 1:500; HPA042781; Sigma), anti-SNAP (WB 1:500; CAB4255, Invitrogen), and anti-WDR44 (WB 1:1000; A301-440A, Bethyl). Membranes were incubated with secondary HRP-labelled antibodies anti-rabbit-IgG (WB 1:5000; 7074S, CST) or anti-mouse-IgG (WB 1:5000; 7076S, CST) in PBST at room temperature for 60 minutes. HRP substrates were used according to the manufacturer’s instructions (170-5060, BioRad; 34096, Thermo Scientific). Immunoblots were imaged using Molecular Imager ChemiDoc XRS+ System (Bio-Rad) and protein bands were quantified using ImageLab (version 6.0.1, BioRad).

### Statistical analyses and data presentation

GraphPad Prism 9 was used for statistical analyses and presentation of quantitative data. The probability values by one way ANOVA test are following: * p<0.05, ** p<0.01, *** p<0.001, **** p<0.0001. Fluorescence microscopy images were processed using ImageJ (Schindelin et al., 2012).

## Acknowledgments

We are grateful to Steve Caplan (University of Nebraska), Derek Toomre (Yale University), Felix Rivera-Molina (Yale University) and other colleagues for discussions and critical reading of the manuscript. We thank Julia von Blume (Yale University), Pietro De Camilli (Yale University), David Calderwood (Yale University) and Shawn Ferguson (Yale University) for sharing reagents and facilities. The authors declare no conflicts of interest. This work was supported by funds from the National Institute of General Medical Sciences of the National Institutes of Health under award numbers GM144096 (CGB) and GM095766-08S1 (CGB).

## Figure Legends

**Figure S1.**
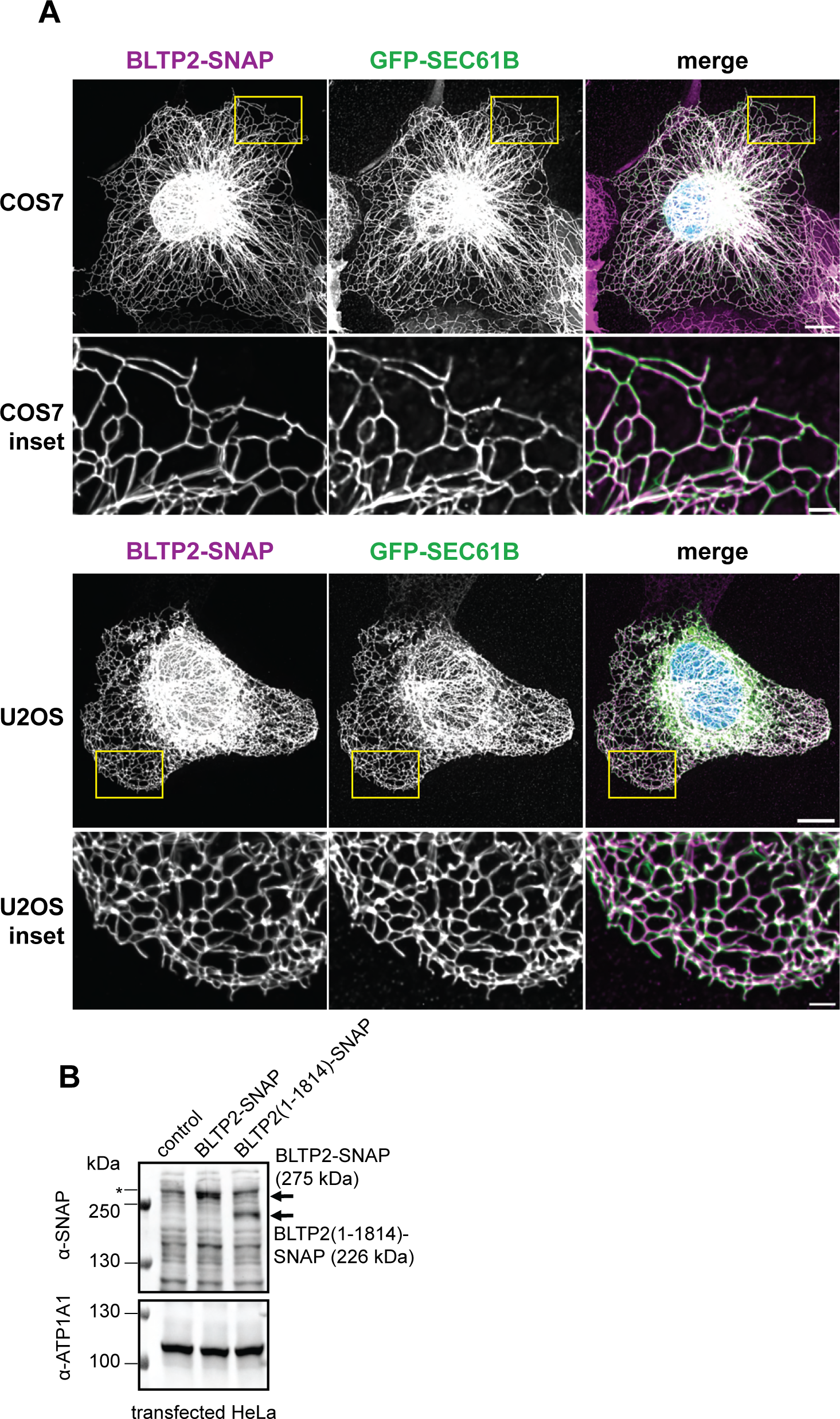
BLTP2 localizes to the ER in cells lacking enlarged tubular endosome. **(A)** COS7 and U2OS cells were transfected with BLTP2-SNAP and GFP-Sec61B, incubate with a SNAP ligand and visualized by SoRa live microscopy. Scale bars: 10 µm; scale bars of insets: 2 μm. **(B)** BLTP2-SNAP and BLTP2ΔC-SNAP are expressed at expected sizes. A representative immunoblot shows the BLTP2-SNAP band (middle) next to the BLTP2ΔC-SNAP band (right). The “control” lane (left) contains lysate from non-transfected cells and the “BLTP2-SNAP” and “BLTP2ΔC-SNAP” lanes contains lysate from the cells transfected with relevant plasmids. The asterisk (*) indicates a non-specific band.

**Figure S2.**
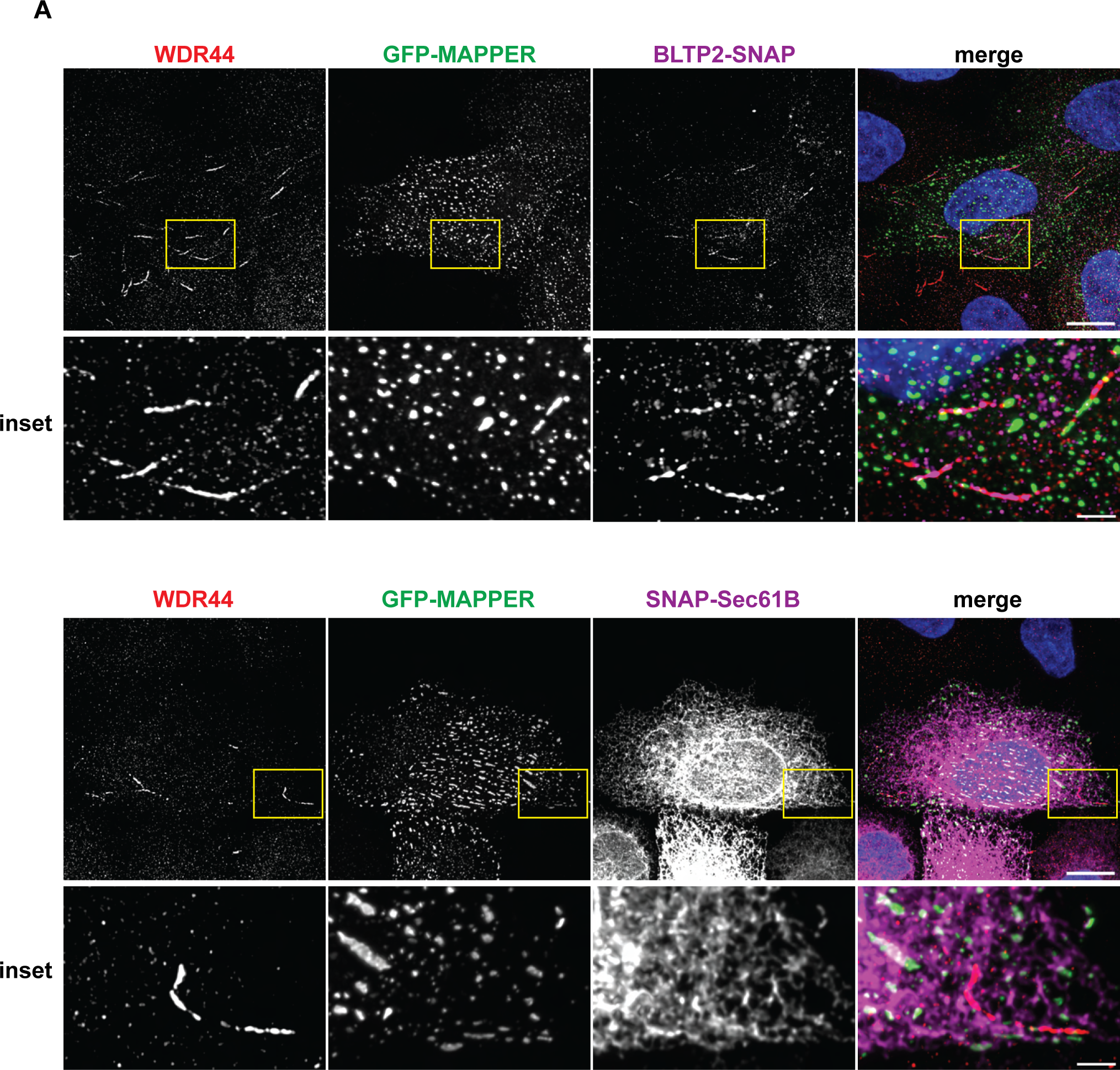
BLTP2 is not enriched at ER-PM contact site in HeLa cells. **(A)** HeLa cells were transfected with GFP-MAPPER and BLTP2-SNAP or SNAP-Sec61B. Cells were first incubated with a SNAP-Cell ligand and then fixed and immunostained using α-WDR44 antibody. Scale bars: 10 µm; scale bars of insets: 2 μm.

**Figure S3.**
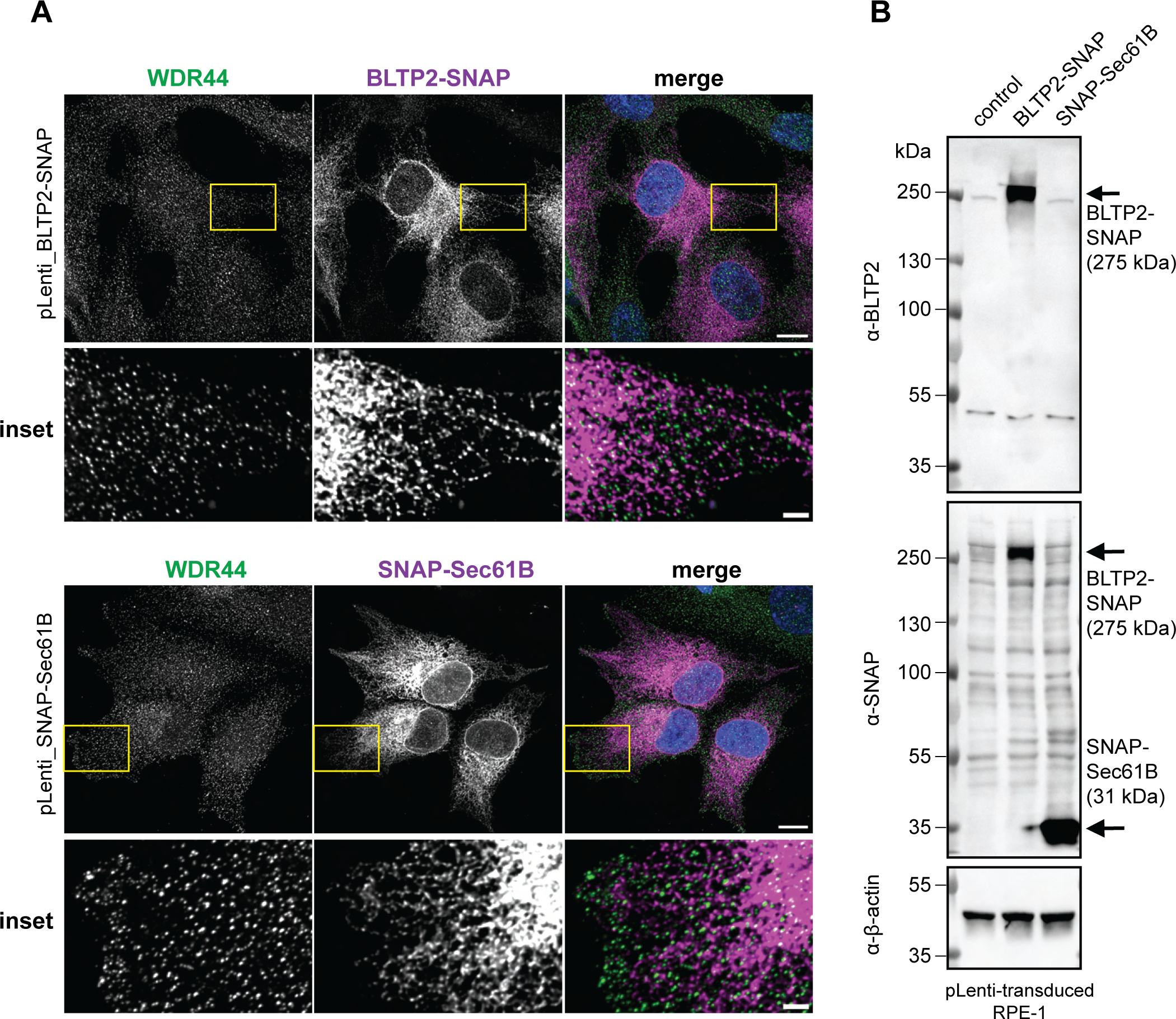
RPE-1 cells do not possess WDR44 tubules. **(A)** RPE-1 cells do not possess WDR44- and BLTP2-associated tubules (≥2 µm). Cells were transduced with pLenti-CMV-Neo-DEST_BLTP2-SNAP or pLenti-CMV-Neo-DEST_SNAP-Sec61B were first incubated with a SNAP-Cell ligand and then fixed and immunostained using α-WDR44 antibody. Scale bars: 10 µm; scale bars of insets: 2 μm. **(B)** RPE-1 cells transduced with pLenti-CMV-Neo-DEST_BLTP2-SNAP express BLTP2-SNAP at expected size (275 kDa).

